## Supplementary information for "Psychedelic 5-HT2A agonist increases spontaneous and evoked 5-Hz oscillations in visual and retrosplenial cortex"

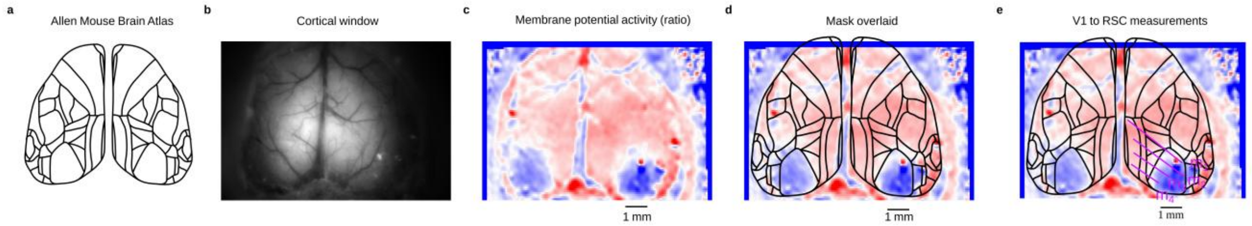


**Supplementary Figure 1: Identification of wide-field cortical regions. a**: Regions of the cortical surface was recreated using the Allen Mouse Brain Atlas (brain-map.org). **b**: An example of the cortical window field of view. **c**: The mean (N=30 trials) hyperpolarisation (identified using the local minima following visually evoked depolarisation) of the membrane potential activity within the cortical window of **b** following a visual stimulus (100% horizontal moving grating for 0.2s). **d**: Manually aligned cortical regions shown in **a** to the regions to the depolarisation in V1, then scaled to the experimental magnification factor. **e**, Estimated distances from V1 to RSC (m1 = 3.166 mm, m2 = 2.884 mm, m3 = 1.968 mm, m4 = 1.486 mm), measured using optical imaging with a calibrated millimeter scale (measurement calibration performed in ImageJ).

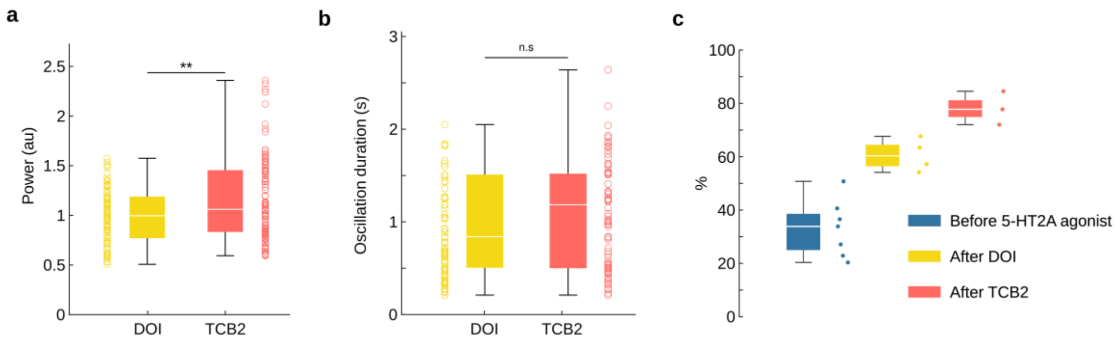


**Supplementary Figure 2: Drug dependency of evoked oscillations in V1. a**, power of oscillations after treatment of DOI (yellow) and TCB2 (red), comparing the individual trials (**, p-value=1.09e-03, Welch's t-test). Data based upon N = 7 experiments in 5 mice; $N_{DOI}$=4; $N_{TCB2}$=3). **b**, same as ***a***, for duration of oscillations across individual trials (n.s, p-value=0.0796, Welch's t-test). **c,** TCB2 induced a more pronounced effect in the probability of 5-Hz oscillations occurring in V1 (oscillations detected using FFT analysis).

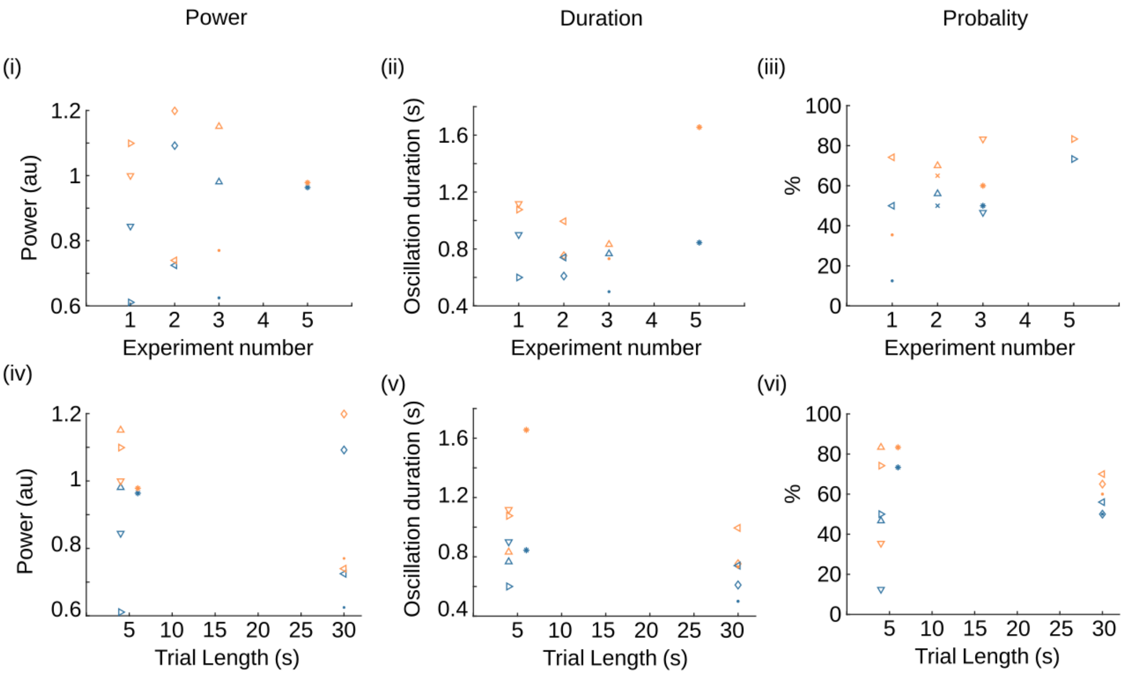


**Supplementary Figure 3: Independence of modulation for evoked 5-Hz oscillations in V1. i-iii**, shows the independence of power, duration and probability against trial length. **iv-vi**, same as **i-iii** for the number of similar experiments the mouse had carried out at each data point.

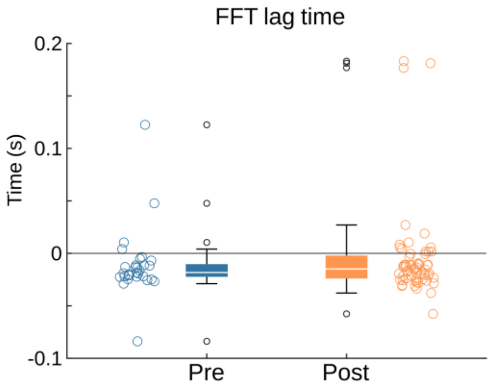


**Supplementary Figure 4: Secondary method of calculating lag time of 5-Hz oscillations between V1 and RSC.** The lag time between signals in V1 and RSC across single trials, using FFT analysis to calculate the phase difference and subsequently the lag time. Showing a median lag of 18.22ms (before, blue) and -14.85ms (after, orange).

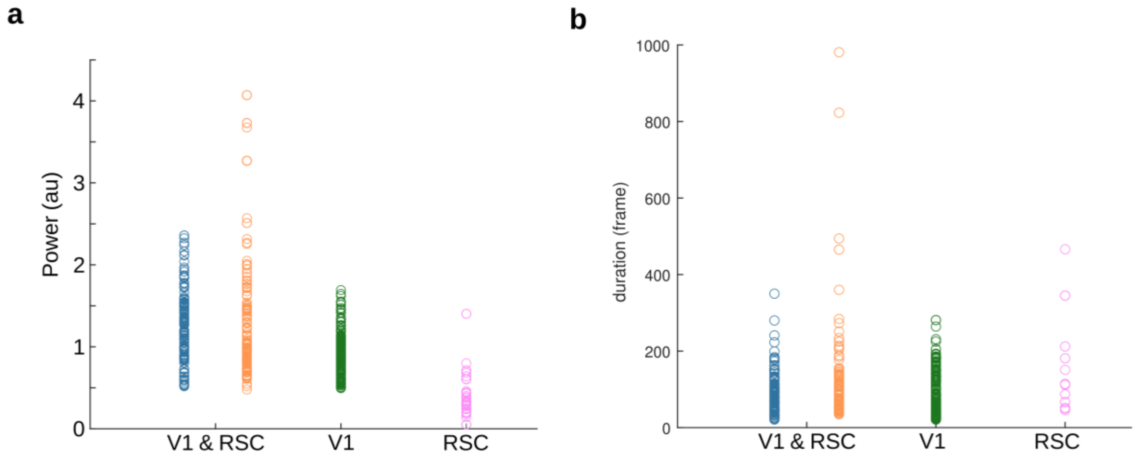


**Supplementary Figure 5: Power and duration for region specific evoked oscillations. a**, the power for evoked oscillations co-occurring in V1 and RSC, V1 only and RSC only. **b**, same as **a**, for duration of evoked oscillations.

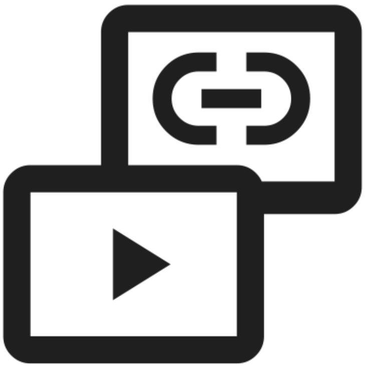

**Supplementary video 1: Visualisation of the evoked 5-Hz oscillation dynamics in V1 and RSC.** Mean activity across trials (N=30, with and without oscillations) based on the experiment shown in **Fig. 3a-b**, before (left) and after (right) injection of a 5-HT2A agonist. Above, showing the spatiotemporal dynamics of the cortex-wide voltage activity (regions outlined using the Allen Mouse Brain Atlas, brain-map.org) and the spatial mean (below) across V1 (green) and RSC (pink). (video file: Supplementary_video_1.avi)
